# Apomixis and genetic background affect distinct traits in *Hieracium pilosella* L. grown under competition

**DOI:** 10.1101/2020.12.30.424832

**Authors:** Christian Sailer, Simone Tiberi, Bernhard Schmid, Jürg Stöcklin, Ueli Grossniklaus

## Abstract

**Background:** Apomixis, the asexual reproduction through seeds, occurs in over 40 plant families and avoids the hidden cost of sex. Apomictic plants are thought to have an advantage in sparse populations and when colonizing new areas but may have a disadvantage in changing environments because they propagate via fixed genotypes.

In this study, we separated the influences of different genetic backgrounds (potentially reflecting local adaptation) from those of the mode of reproduction, i.e., sexual vs. apomictic, on nine fitness-related traits in *Hieracium pilosella* L. We aimed to test whether apomixis *per se* may provide a fitness advantage in different competition environments in a common garden setting.

**Results:** To separate the effects of genetic background from those of reproductive mode, we generated five families of apomictic and sexual full siblings by crossing two paternal with four maternal parents. Under competition, apomictic plants showed reproductive assurance (probability of seeding, fertility), while offspring of sexual plants with the same genetic background had a higher germination rate. Sexual plants grew better (biomass) than apomictic plants in the presence of grass as a competitor but apomictic plants spread further vegetatively (maximum stolon length) when their competitors were sexual plants of the same species. Furthermore, genetic background as represented by the five full-sibling families influenced maximum stolon length, the number of seeds, and total fitness. Under competition with grass, genetic background influenced fecundity, the number of seeds, and germination rate.

**Conclusions:** Our results suggest that both the mode of reproduction as well as the genetic background affect the success of *H. pilosella* in competitive environments. Total fitness, the most relevant trait for adaptation, was only affected by the genetic background. However, we also show for the first time that apomixis *per se* has effects on fitness-related traits that are not confounded by — and thus independent of — the genetic background.

## Background

Apomixis in higher plants is viewed as a deregulation of sexual processes in space and time [1–4]. In apomicts, three steps of the sexual process are altered: (i) meiosis is aberrant or omitted (apomeiosis), leading to the production of unreduced gametes; (ii) embryogenesis is initiated without fertilization of the egg cell (parthenogenesis); and (iii) endosperm development can be autonomous (no fertilization of the central cell) or pseudogamous, i.e., require fertilization [5] with certain developmental adaptations that ensure normal seed development [4]. In combination, the three elements of apomixis result in the production of maternal clonal offspring [1,5], a trait of evolutionary, ecological, and agronomic importance [6,7].

Apomixis with autonomous endosperm development results in reproductive assurance, because these lineages do not require pollination by selfing or crossing [8–11]. Such apomictic plants are thought to have an advantage in sparse populations [9] and can even found new populations developing from a single individual [Baker’s law, 12]. Typically, apomixis only affects female gametophyte development while male gametophyte development is normal such that apomictic plants can outcross via pollen [1]. Furthermore, apomixis is a quantitative, facultative trait [13–15]. As a result of its facultative nature and normal male function, apomixis is not a dead end of evolution [16]. Nonetheless, the clonal nature of reproduction in apomicts could result in a reduced adaptive potential under environmental change [17–19], an issue that has not yet been experimentally addressed. At the same time, apomixis allows the fixation of genotypes that are well adapted to current environmental conditions, providing an advantage for population expansion.

In the case of the autonomous aposporous apomict *Hieracium pilosella* L., which is native to central Europe, apomixis has likely helped the successful invasion of new geographic ranges with similar environmental conditions as in the home range, e.g., New Zealand [20–23] and Patagonia [24]. Furthermore, in this species apomixis is a facultative, quantitative trait [13] and sexual reproduction can occur in natural populations, enabling the generation of new (facultatively apomictic) genotypes that are adapted to novel environments [25]. In New Zealand, *H. pilosella* was introduced several times and hybridized with the closely related *H. praealtum* [26,27], creating new genotypes that allowed for rapid population expansion.

In a previous study [28], we found that invasive, apomictic pentaploid genotypes of *Hieracium pilosella* L. from New Zealand (aP5) had a higher competitiveness compared with sexual tetraploid genotypes (sP4). On the one hand, the apomictic plants had higher biomass and competitiveness due to higher ploidy, on the other hand, not all sexual genotypes had the same performance. This raises the question to what extent the mode of reproduction (apomixis vs. sexual) *per se* influences competitiveness, independently of the rest of the genome (genetic background). To investigate this question, we generated several F1 hybrid families of hexaploid apomictic (aP6) and sexual plants (sP6) with highly similar genetic background (full-siblings). Individuals of a full-sibling family are as genetically close as possible in an obligately outcrossing species. We used these families to separate the influence of the genetic background from that of the mode of reproduction on plant performance in a competitive environment. In this study, the families represent distinct genetic backgrounds independent of the reproductive mode. We investigated these apomictic and sexual siblings in the same setting as in a previous study [28] in order to be able to compare the results of the two analyses. We measured three vegetative and six generative fitness-related traits (Table 1). In particular, we tested for which traits competitiveness differed (i) between apomictic plants and sexual plants, and (ii) between different genetic backgrounds corresponding to the parental genotypes used in the crosses.

**Table 1.**
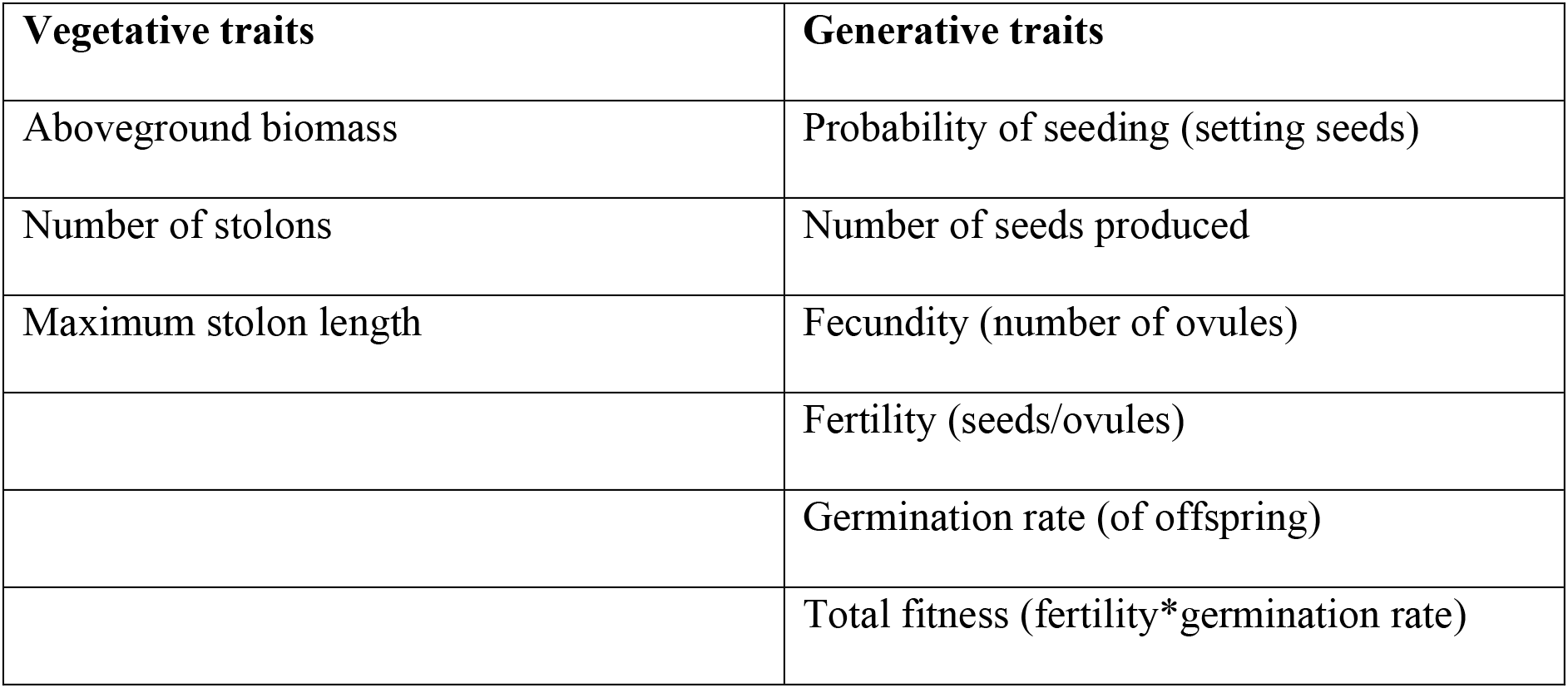
Measured and computed fitness-related traits

| **Vegetative traits** | **Generative traits** |
| --- | --- |
| Aboveground biomass | Probability of seeding (setting seeds) |
| Number of stolons | Number of seeds produced |
| Maximum stolon length | Fecundity (number of ovules) |
|  | Fertility (seeds/ovules) |
|  | Germination rate (of offspring) |
|  | Total fitness (fertility*germination rate) |

Apomictic and sexual plants responded differently to competition with regard to biomass, maximum stolon length, probability of seeding, fertility, and germination rate of the offspring. The families (genetic background of the parental genotypes) influenced stolon length, the number of seeds, germination rate, and total fitness. In presence of within-species competition (neighbors of different reproductive mode), sexual plants performed better in terms of germination rate and fitness of seed-producing plants, while in between-species competition (presence of grass), apomictic plants performed better in terms of fertility. Furthermore, under between-species competition, fecundity, the number of seeds, and the germination rate of the offspring varied among families. Our findings indicate that, across all traits, both the mode of reproduction and the genetic background affect fitness-related traits and, thus, competitiveness of *H. pilosella*. We found that reproductive mode affects some traits independent of the genetic background (biomass, probability of seeding, fertility). The number of seeds and total fitness were affected by genetic background (family) only.

## Results

### Genetic background, mode of reproduction, and interspecific competition have distinct effects on measured traits

We found that overall competition with grass reduced biomass (F_1, 25.4_ = 13.42, P = 0.0011, Figure 1a, Supplementary Table 1a), fecundity (F_1, 21.5_ = 4.14, P = 0.0544, Figure 1b, Supplementary Table 1e), the number of seeds (F_1, 24.9_ = 6.85, P = 0.0148, Figure 1c, Supplementary Table 1f), and fertility (F_1, 22.3_ = 5.95, P = 0.0231, Figure 1d, Supplementary Table 1g).

**Figure 1:**
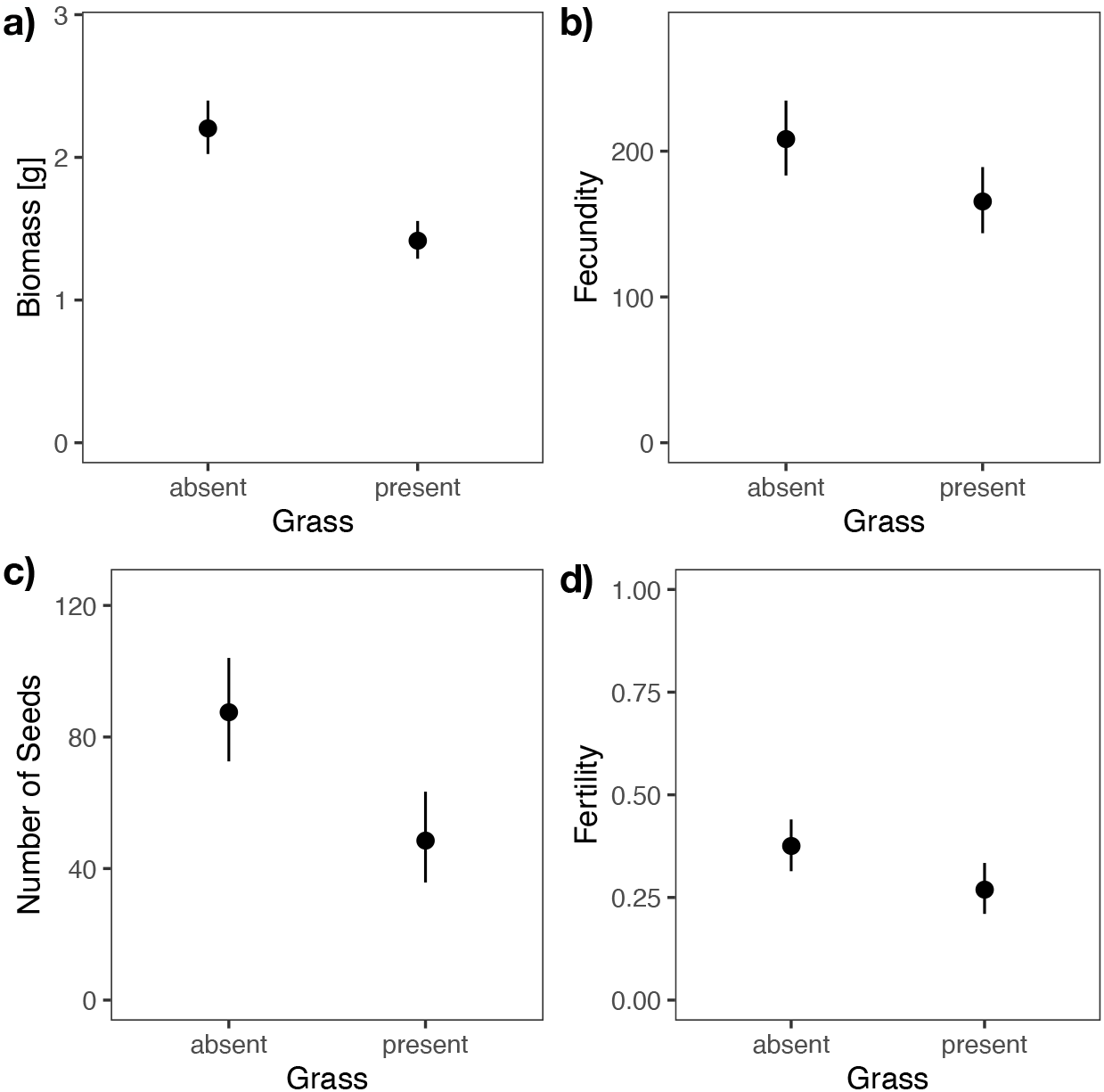
Effects of competition with grass on four traits. **a)** Biomass, **b)** Fecundity, **c)** Number of seeds (produced), **d)** Fertility. Dots are point estimates from the model and error bars show ± 1 standard error.

The five families differed overall in their maximum stolon length (F_4, 10.7_ = 5.41, P = 0.0123, Figure 2a, Supplementary Table 1b), the number of seeds (F_4, 8.9_ = 4.15, P = 0.0357, Figure 2b, Supplementary Table 1f), germination rate (F_4, 9.4_ = 4.35, P = 0.0297, Figure 2c, Supplementary Table 1h), and total fitness (F_4, 8.0_ = 4.38, P = 0.0362, Figure 2d, Supplementary Table 1i).

**Figure 2:**
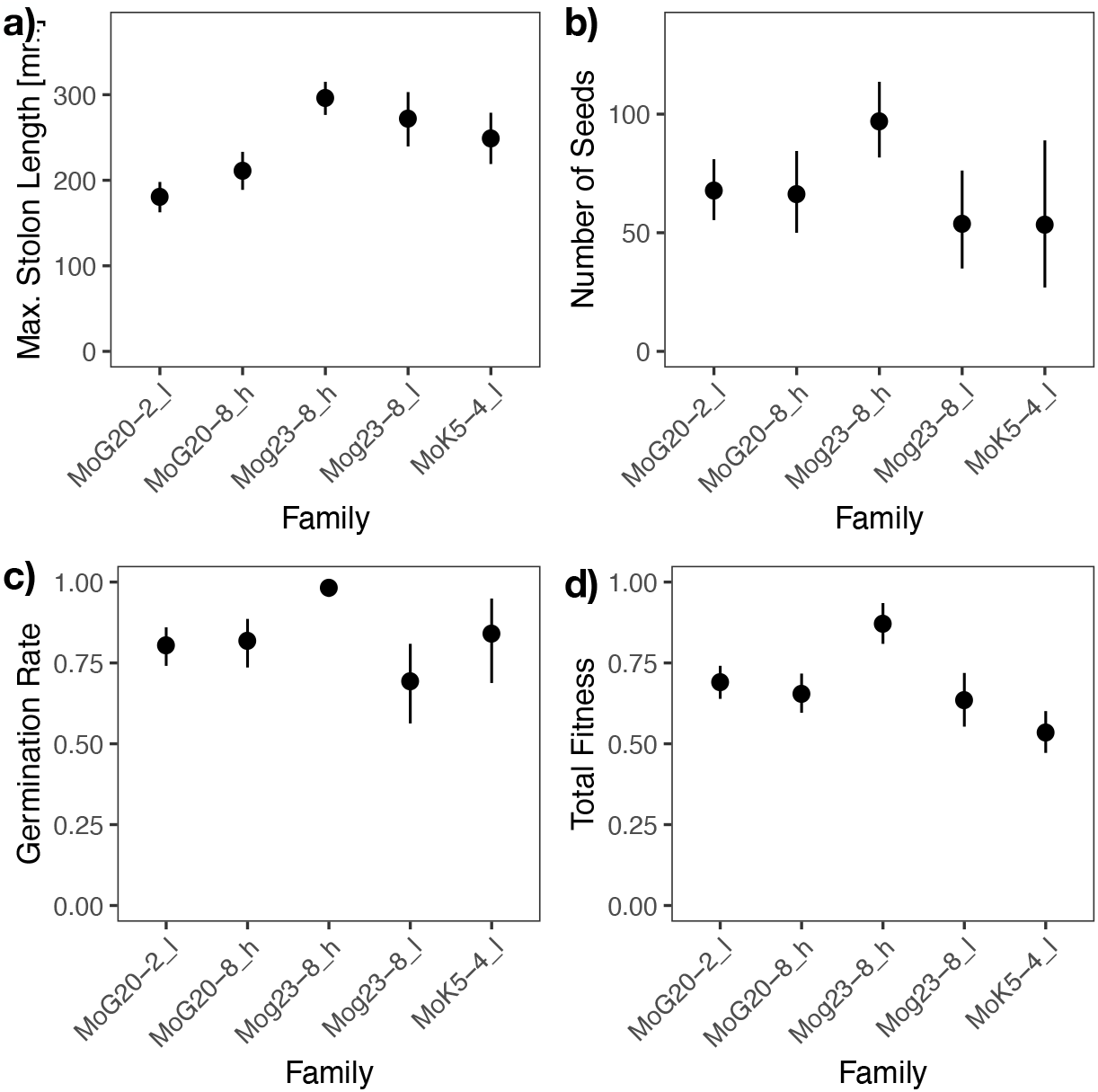
Differences among five full-sibling families. **a)** Maximum stolon length, **b)** Number of seeds (produced), **c)** Germination rate (of offspring), **d)** Total fitness. Dots are point estimates from the model and error bars show ± 1 standard error. ‘l’ and ‘h’ in the family designation indicate low and high expressivity of apomixis in the father.

Moreover, we found that apomictic plants had, overall, a higher probability of seeding (F_1, 12.3_ = 5.42, P = 0.0377, Figure 3a, Supplementary Table 1d) and a lower germination rate of their offspring (F_1, 10.9_ = 11.26, P = 0.0065, Figure 3b, Supplementary Table 1h) than sexual plants.

**Figure 3:**
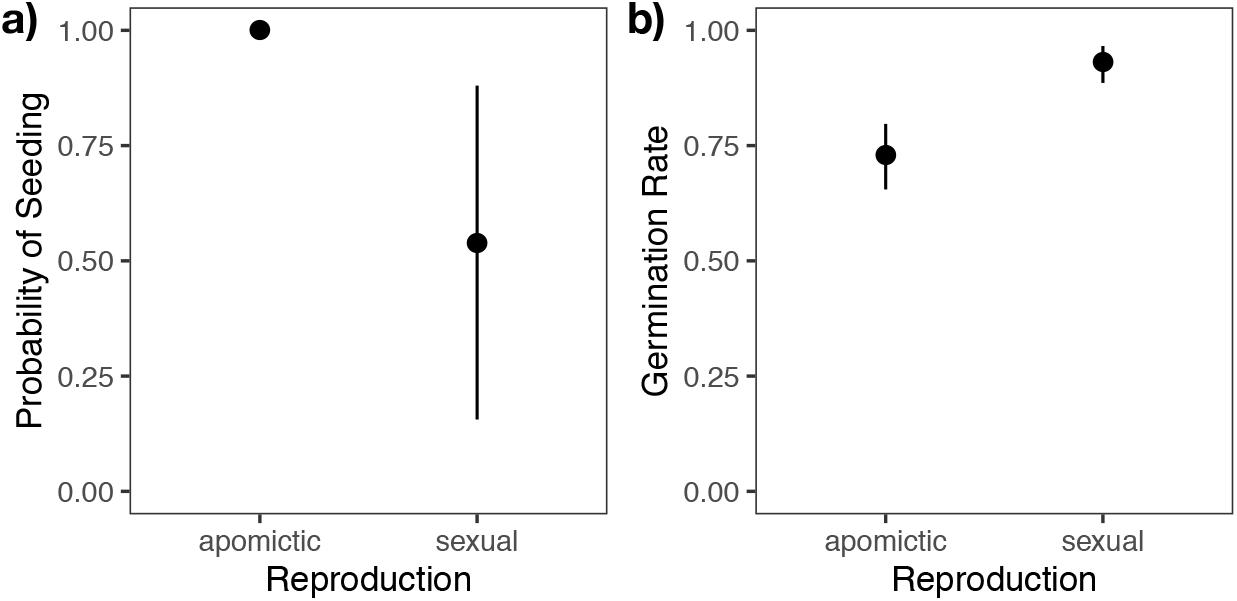
Differences between apomictic and sexual siblings. **a)** Probability of seedings (= setting seeds), **b)** Germination rate (of offspring). Dots ae point estimates from the model and error bars show ± 1 standard error.

### Genetic background affects the response to competition

We found that families did not differ in their response to the presence of grass in terms of fecundity (F_4, 64.0_ = 1.75, P = 0.1498, Supplementary Table 1e). However, when we considered the contrast between high- and low-apomixis fathers (paternal half-sibling families), this effect became significant (P = 0.043, Figure 4a). The difference between plants with a low- or a high-apomixis father did not significantly affect any other trait tested. Furthermore, families showed different responses to competition with grass in terms of the number of seeds (F_4, 58.8_ = 3.82, P = 0.0080, Figure 4b, Supplementary Table 1f) and germination rate (F_4, 52.9_ = 4.04, P = 0.0062, Figure 4c, Supplementary Table 1h).

**Figure 4:**
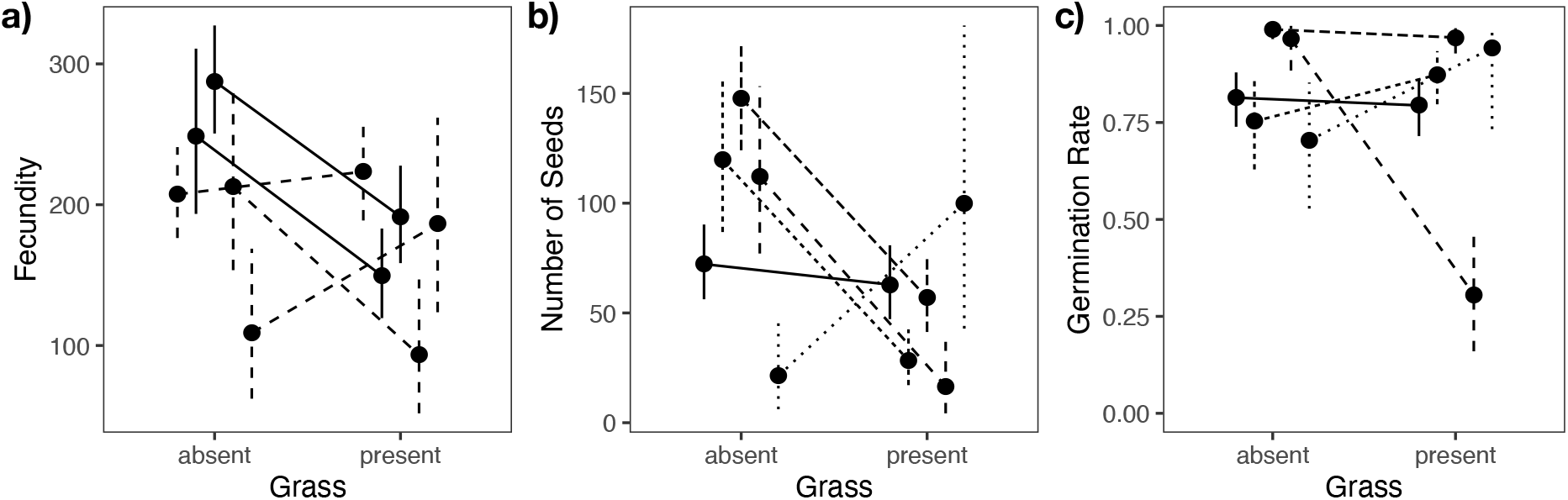
Different response of the five full-sibling families to competition with grass. **a)** Fecundity shows a different response between fathers (paternal half-sibling families), indicated by the two different line types. Solid lines indicate high level and dashed lines low levels of apomixis of the father. **b)** Number of seeds (= seed set), **c)** Germination rate (of offspring). Dots are point estimates from the model and error bars show ± 1 standard error.

### The response to competition depends on reproductive mode

Apomictic and sexual plants showed different competitiveness. The presence of grass reduced the growth of apomictic plants (F_1, 106.4_ = 7.58, P = 0.0067, Figure 5a, Supplementary Table 1a). Furthermore, the presence of grass reduced fertility (F_1, 69.7_ = 13.43, P = 0.0005, Figure 5b, Supplementary Table 1g) when the plants were sexual but not when they were apomictic, indicating that apomictic plants were weaker vegetative but better generative between-species competitors.

**Figure 5:**
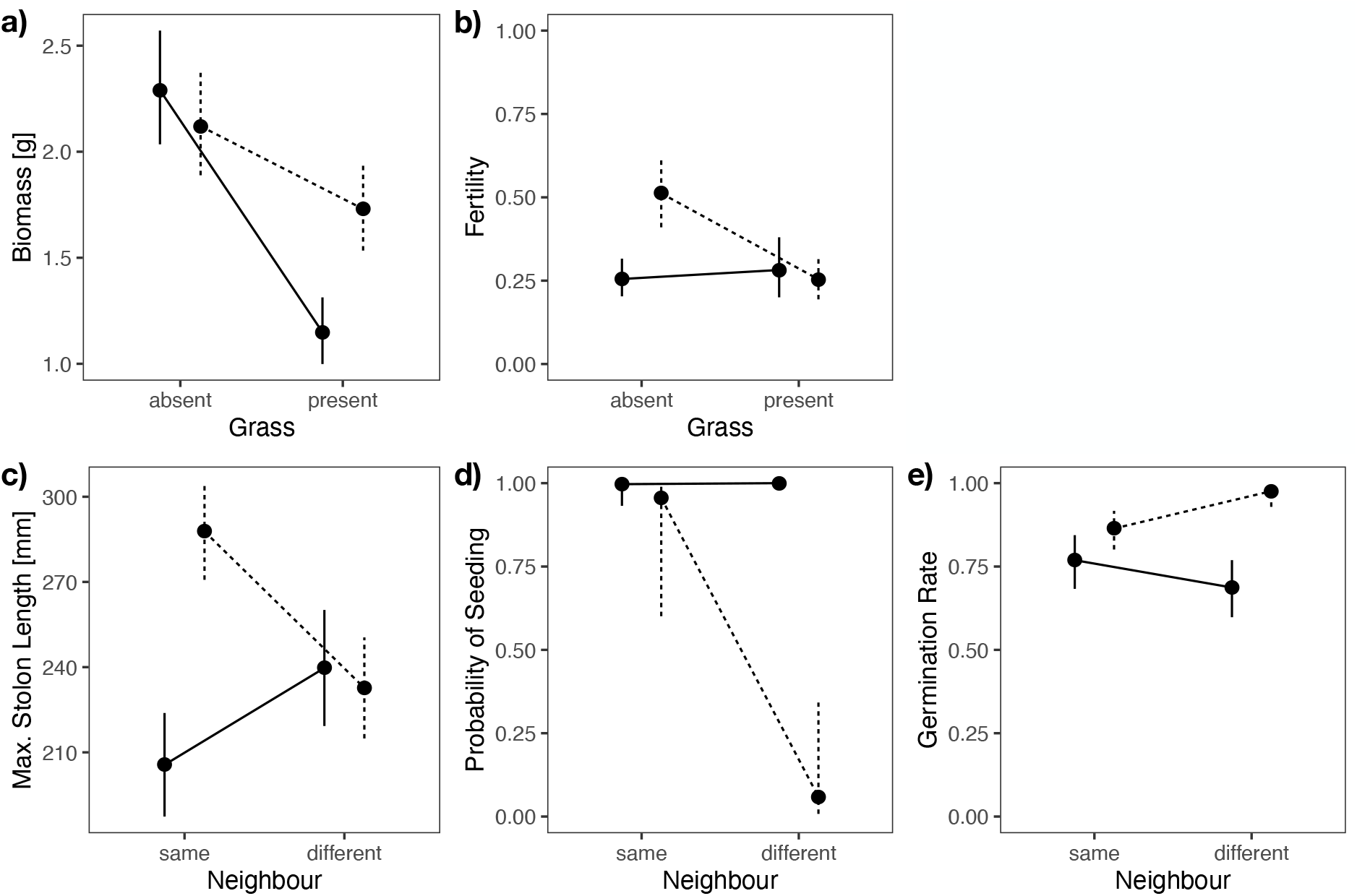
Apomictic and sexual plants react differently to competition with grass (a-b) and to within-species competition with plants of the other mode of reproduction (c-e). **a)** Biomass, **b)** Fertility, **c)** Maximum stolon length, **d)** Probability of seeding, and **e)** Germination rate (of offspring). Dots are point estimates from the model and error bars show ± 1 standard error. Solid lines represent apomictic and dashed lines sexual plants, respectively.

Having a neighbor of different reproductive mode (competition within *Hieracium*) increased the maximum stolon length per plant in apomictic plants while it reduced it in sexual plants (F_1, 85.3_ = 10.46, P = 0.0017, Figure 5c, Supplementary Table 1b). This indicates that, with respect to vegetative dispersal, apomictic plants show a better within-species competitiveness. Furthermore, the probability of seeding in apomictic plants was not affected by competition with sexual plants (F_1, 62.2_ = 4.96, P = 0.0295, Figure 5d, Supplementary Table 1d). However, offspring from apomictic plants had a lower germination rate than offspring from sexual plants (F_1, 40.1_ = 6.44, P = 0.0151, Figure 5e, Supplementary Table 1h). These findings suggest that, while apomictic plants set seed irrespective of competition, sexual plants were better within-species competitors.

### Effects of combined competition treatments on spread via stolons

Overall, the combination of within- and between-species competition as well as no competition increased both stolon length and the number of stolons, whereas shorter and fewer stolons were produced with only one type of competition (stolon length: F_1, 21.2_ = 35.57, P < 0.0001, Figure 6a, Supplementary Table 1b; number of stolons: F_1, 25.6_ = 6.79, P = 0.0150, Figure 6b, Supplementary Table 1c). In other words, for vegetative spread and propagation having more competition is as beneficial as having no competition with plants of the other reproductive mode or with grass.

**Figure 6:**
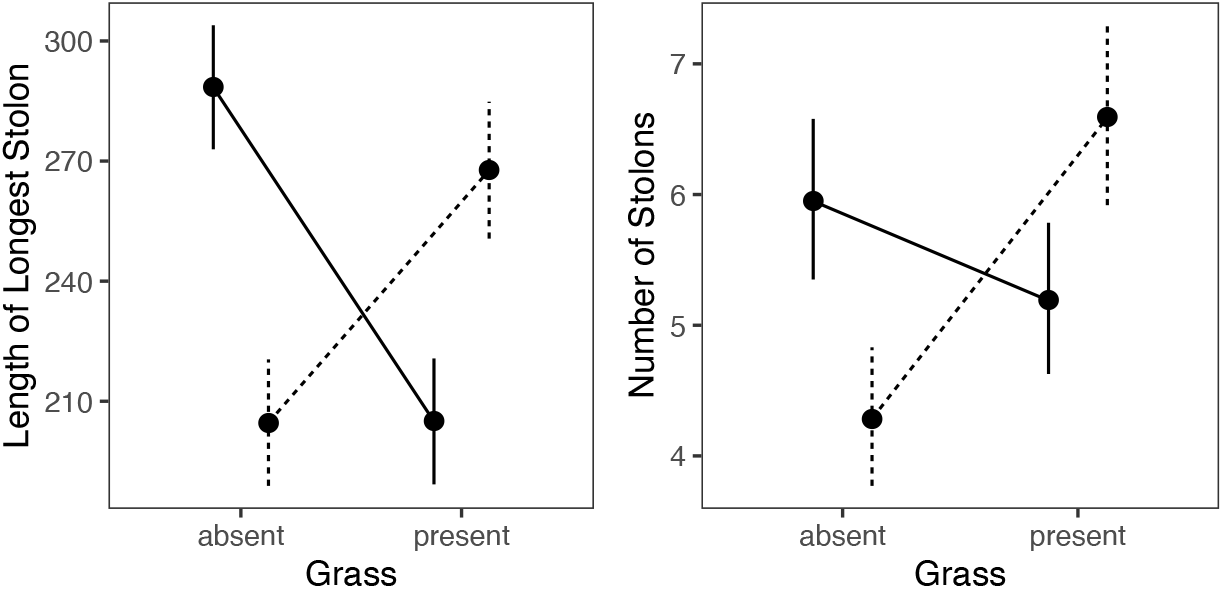
Interactions between within-species competition and competition with grass. **a)** Length of longest stolon, **b)** Number of stolons. Dots are point estimates from the model and error bars show ± 1 standard error. Solid lines for no within-species competition, dashed lines for competition against plants with the other mode of reproduction.

## Discussion

With this experiment, we aimed to investigate whether the mode of reproduction *per se* influences competitiveness and to which extent the genetic background is of importance for differences in performance of *Hieracium pilosella* lines. In our previous study, we compared pentaploid apomictic with sexual plants but, as the latter were sterile, we could not measure generative traits [28]. Here, we investigated hexaploid apomictic and sexual plants as they occur in natural populations [25,29].

### Effects of competition on vegetative traits

In concert with our previous work, we detected effects of competition on biomass production in apomictic and sexual full-sibling families. However, in the present study, we found that grass affected apomictic plants more strongly, while previously we did not detect this difference in competitiveness. Instead, we had found that within-species competition favored apomictic plants. Together with the ploidy effect and different genetic backgrounds between apomictic and sexual plants (apomictic and sexual plants belonged to unrelated families) in our previous study, we can now conclude that the apomictic families used in the previous study were better growth competitors, i.e., that this effect was likely due to their genetic background. Furthermore, it explains why we had not detected different competitiveness with grass in the previous study. Here, because of the use of full-sibling families, we can exclude effects of the genetic background between apomictic and sexual plants, which may explain why we did not detect different within-species competitiveness anymore but still different between-species competitiveness. Taking the results of both studies together, we suggest that apomictic plants are weaker between-species growth competitors than sexual plants are. This finding suggests that the apomictic genotypes that are invasive in New Zealand [21,26,27] were selected for fast growth under competition. The fact that the vast majority of the genotypes in New Zealand are pentaploid [30,31] leaves apomixis as their only option to reproduce via seeds (reproductive assurance) [16].

In terms of vegetative propagation via stolons (stolon count), we did not detect any differences between apomictic and sexual plants using our full-sibling families. In the previous study, we found that apomictic plants produced fewer stolons in competition with grass. We now tentatively attribute this previously observed difference to genetic background (family) and not reproductive mode. Hence, we conclude that vegetative propagation is likely independent of the reproductive mode.

In terms of vegetative spread, we previously found that sexual plants were less affected by grass than apomictic plants were. Using our full-sibling families, this effect disappeared. Hence, we can attribute the previously found difference to differences in genetic background (family) rather than reproductive mode. Instead, we found that sexual plants produced longer stolons when grown together with their apomictic full siblings, suggesting an association between mode of reproduction and vegetative spread. However, full-sibling families differed among each other, indicating that vegetative spread is determined by a combination of the mode of reproduction and genetic background.

### Generative traits depend on the mode of reproduction

In terms of germination rates of offspring, sexual plants outperformed apomictic plants in general and under within-species competition. We found support for reproductive assurance [16] in apomictic plants as they were more likely to set seed in general and in competition with sexual plants. Moreover, in contrast to sexual plants, apomictic plants did not show reduced fertility in the presence of grass, further supporting reproductive assurance as an advantage of apomictic lineages. However, reproductive assurance seemed to be counteracted by the higher germination rate of seeds from sexual plants. When we summarized fertility and germination rate into fitness, we found no difference between the two modes of reproduction, suggesting that apomixis is not a dead end of evolution [16]. It is important to mention that, when considering only plants that had set seeds in the estimation of fitness, apomictic plants were superior. However, this effect is a direct consequence of reproductive assurance via apomixis. Hence, in environments where setting seeds is the most relevant trait, like in sparse population densities or when there is a lack of pollinators, apomictic plants would have a clear competitive advantage [9]. However, in the case of *H. pilosella*, this prediction was not supported by characterizing a natural population in the Swiss alps [25].

Overall, the reproductive mode affected competitiveness in a variety of traits and at different levels of competition. However, for about half of the traits analyzed, apomictic plants were better competitors while for the other half, sexual plants were superior. Hence, we could not assign a clear competitive advantage to either mode of reproduction.

### The role of the genetic background

To separate the influence of the genetic background from the effects of the mode of reproduction, we used a total of five full-sibling families. Using such newly generated F1 full-sibling families removed potential biases that may stem from local adaptation in the parental lineages [32], because both apomictic and sexual plants were equally likely to inherit the locally adapted alleles. This allowed us to interpret the effects of the genetic background without local-adaptation bias and disentangled from the mode of reproduction.

The genetic background (full-sibling family) was relevant for between-species competitiveness with grass in three reproductive traits (fecundity, number of seeds, germination rate). In general, families differed in the maximum stolon length, the number of seeds, germination rate, and, importantly, total fitness. No other tested factor was relevant for differences in total fitness. With full-sibling families representing genetic diversity in our experiment, we can once again point out the relevance of variation and diversity for adaptation and evolution, as they are the basis on which natural and artificial selection can act upon [33]. Our results thus reflect differences in performance that are based on genetic diversity between the five families that we used in this experiment, which was independent of the mode of reproduction.

## Conclusions

We found that both the family, i.e., genetic background, and mode of reproduction affected fitness-related traits in the apomictic species *Hieracium pilosella*, particularly in their response to different competitive environments. To the best of our knowledge, the present study is the first experimental demonstration that, within a given generation, the mode of reproduction — independent of genetic background — is of relevance for the success of *H. pilosella* plants in different environments. The mode of reproduction is important for traits that are a consequence of reproductive assurance (probability of setting seed and fertility). However, when considering future generations, adaptation and the formation of new genotypes, i.e., total fitness that includes all individuals of a population, is the more relevant trait. We found that only genetic background, i.e. differences among full-sibling families, affected total fitness. Unless reproductive assurance is of utmost importance for reproductive success in a given environment, genetic diversity is the most relevant factor for reproductive success in *H. pilosella* L.

## Materials and Methods

### Plant Material

*Hieracium pilosella* L. is a self-incompatible, perennial, monocarpic, herbaceous species including sexual and apomictic lineages, which can occur at different ploidy levels [29]. Plants can reproduce vegetatively via aboveground stolons. Apomictic lineages are of the autonomous apospory type [11] and apomictic plants can outcross via pollen.

We isolated four sexual hexaploid genotypes (sP6, **s**exual **P**ilosella **6**-ploid) from a population at the forefield of the Morteratsch glacier in the Upper Engadin in Switzerland (MoK5-4, MoG20-2, MoG20-8, MoG23-8, latitude: 46.43420, longitude: 9.93537) and two apomictic hexaploid genotypes (aP6, **a**pomictic **P**ilosella **6**-ploid) from two populations in New Zealand (genotype LaP1, Lake Pukaki, latitude: –44.15848, longitude: 170.22020 and genotype MwR1, Molesworth Road, latitude: –42.00933, longitude: 172.95406). Genotype LaP1 had low apomictic fertility (low), while genotype MwR1 had high apomictic fertility (high). We crossed both apomictic genotypes as pollen donors (fathers) to four sexual maternal genotypes and obtained five full-sibling families with a father either with a low or a high expressivity of apomixis. The F1 plants resulting from these crosses were grown in the greenhouse and tested for apomixis by decapitation [34]. In this way, we generated and identified apomictic and sexual full-sibling families (highly similar genetic background), which were then propagated vegetatively to generate eight clonal replicates per plant.

### Experimental Design

We used the same experimental design as Sailer and colleagues [28]. To define the genotype combinations for the experiment, we randomly selected apomictic and sexual plant pairs from each of the full-sibling families. Since not all crosses to generate the families were equally successful, we randomly combined pairs that had fathers with low or high expressivity of apomixis to ensure a controlled genetic diversity within each replicate and a random distribution of families across the entire experiment. Furthermore, this design allowed for an additional comparison at the level of expressivity of apomixis of the father. That is, pairs sired by the high-apomixis father (Ah – Sh: **A**pomictic, **h**igh-apomixis father – **S**exual, **h**igh-apomixis father) could be compared with pairs sired by the low-apomixis father (Al – Sl: **A**pomictic, **l**ow-apomixis father – **S**exual, **l**ow-apomixis father). This combination of apomictic and sexual full-sibling families from fathers with different expressivity of apomixis was defined for each replicate. ‘High-apomixis-father’ and ‘low-apomixis-father’ plants were always grown together in the same boxes (two different families per box). Apomictic and sexual siblings (two levels of reproductive mode) were grown either alone or together with the other reproductive type (within-species competition), and with or without the grass *Bromus erectus* Huds. (between-species competition). *Bromus erectus* served as between-species competitor because the two species often co-occur in nature. The three treatments mentioned above (reproductive mode, competition by grass, within-species competition) resulted in eight different treatment combinations at the level of boxes, and a fourth treatment within each box (two families with siblings from either low- or high-apomixis fathers), thereby resulting in 16 treatments (Supplementary figure 1).

The plants grew in a total of forty plastic boxes (Georg Utz AG, Bremgarten, Switzerland) of 40 × 30 cm area and 30 cm depth, with each treatment replicated five times, using plants of different full-sibling families of *H. pilosella.* The bottom of the boxes had holes and was covered with a 2 cm thick drainage mat to prevent root rotting in standing water. The boxes were covered with mosquito net (Windhager AG, Baar, Switzerland) cages to prevent pollination between experimental units in the common garden. In each box, the positions of four *H. pilosella* plants were fixed in a grid with a total of eight positions (as on a chessboard on squares of the same color). Individuals of the defined genotype combinations were randomly assigned to these positions. For between-species competition four *H. pilosella* plants were grown alternating with four plants of the grass *B. erectus* Huds. from the Swiss Jura Mountains (Otto Hauenstein Samen, Rafz, Switzerland). For within-species competition, sexual and apomictic full-siblings from both the low- and high-apomixis fathers were planted alternately in a ratio of 2 apomictic : 2 sexual plants. To control for potential position effects, the order of the two was switched in every second experimental unit.

### Crosses, Measurements, and Harvest

Plants grew in the common garden of the Department of Plant and Microbial Biology of the University of Zurich, Switzerland, from April 2012 to September 2012. At the day of opening of a capitulum, we measured the diameter of the opened capitulum to a precision of 1 mm. We crossed all capitula of all individuals of one experimental unit (a single box) that were open at the same time with each other by rubbing two capitula together. Crosses were repeated every day until closing of the capitula. At the day of measuring seed set, we harvested the seeds and stored them at 4 °C and 30% relative humidity until use. At the end of August, most plants had set seed and we started harvesting. We cut off the stolons and collected them separately from the plants without roots. To measure biomass, we oven-dried the plant material for 48 h at 80 °C and weighed the dried material to a precision of 0.1 mg.

To determine fecundity and fertility, we counted harvested seeds and empty seed shells. To determine germination rates, we surface-sterilized up to 20 seeds from one randomly chosen capitulum per individual and germinated them in petri dishes on half-strength MS-medium [35] (MS salts; Carolina, Burlington, North Carolina), Sucrose (Applichem, Darmstadt, Germany), and Phytoagar (Gibco BRL, Paisley, Scotland) in a Percival Scientific climatic cabinet (CU-36L6/D, CLF Plant Climatics GmbH, Wertingen, Germany) at a 14 h, 22 °C light and 10 h, 18 °C h dark cycle after 72 h of stratification at 4 °C. We measured germination success every day for 7 days.

### Measured and Computed Traits

We measured the aboveground biomass of each plant, counted the number of stolons, and measured their length. For generative traits, we counted the total number of seeds and empty seed shells. For computed traits, we determined the probability of seeding (plant producing seeds vs. not producing seeds), fecundity (number of seeds plus number of empty seed shells, only seeding plants, equivalent to the number of ovules), fertility (seed set or number of seeds divided by fecundity, only seeding plants), germination rate (number of germinated seeds divided by number of seeds plated, only seeding plants, and total (Darwinian) fitness (product of fertility and germination rate, all plants).

### Statistical Analysis

We fitted a separate model for each trait. Fecundity, stolon count, and number of seeds were square-root transformed and analyzed with linear models that gave better fits than Poisson or negative-binomial models. For seeding and fertility, we used a binomial model with logit link function. For germination rate, we used the angular transformation (arcsin of square root). Total fitness was transformed with the logarithm (log(fitness+0.5)), while aboveground biomass and maximum stolon length were analyzed on their original scale. In all cases, we used mixed-effects models with line (seven crosses, i.e., one full-sib family was replicated in three crosses) and box (n = 40) as random effects. As fixed effects, we fitted at the box level between-species competition (grass present or absent), within-species competition (neighbor of same or different reproductive mode), and their interaction (omitted if P > 0.1). At the individual level, we fitted the mode of reproduction (sexual or apomictic), full-sibling family (genetic background including different fathers), and their interactions with each other and with the competition treatments. Terms were considered significant if P < 0.05 and marginally significant if P < 0.1. We also made contrasts among the five full-sibling families for the two paternal parents (four crosses representing two full-sibling families vs. three crosses representing three full-sibling families). However, these were rarely much larger than the remaining differences among full-sibling families within paternal parents and, therefore, not included in the analysis of variance tables.

All statistical analyses and plots were done in R [36]; mixed effects models were fitted using the asreml R package [37,38], while plots were done in the ggplot2 R package [39].

## List of abbreviations

aP5: apomictic *Pilosella* 5-ploid (pentaploid)
aP6: apomictic *Pilosella* 6-ploid (hexaploid)
sP4: sexual *Pilosella* 4-ploid (tetraploid)
sP6: sexual *Pilosella* 6-ploid (hexaploid)

## Declarations

### Ethics approval and consent to participate

Not applicable.

### Consent for publication

Not applicable.

### Availability of data and materials

All data generated or analysed during this study are included in this published article and its supplementary information files.

### Authors’ contributions

UG initiated the project; UG and JS acquired funding; CS, JS, BS, UG conceived the study; CS and BS designed the experiment; CS performed the experiment; CS, ST, BS analyzed the data; UG supervised the project; CS wrote the original draft; all authors revised and edited the manuscript.

## Acknowledgements

We thank Alain Held for help with harvesting and measurements, Hannes Vogler for careful proofreading of the manuscript, and Christian Frey and Karl Huwiler for help with plant care. We thank Ross Bicknell for collecting and sharing seeds from New Zealand.

