## Supplementary figures and images for "Apomixis and genetic background affect distinct traits in *Hieracium pilosella* L. grown under competition"

### Supplementary figure

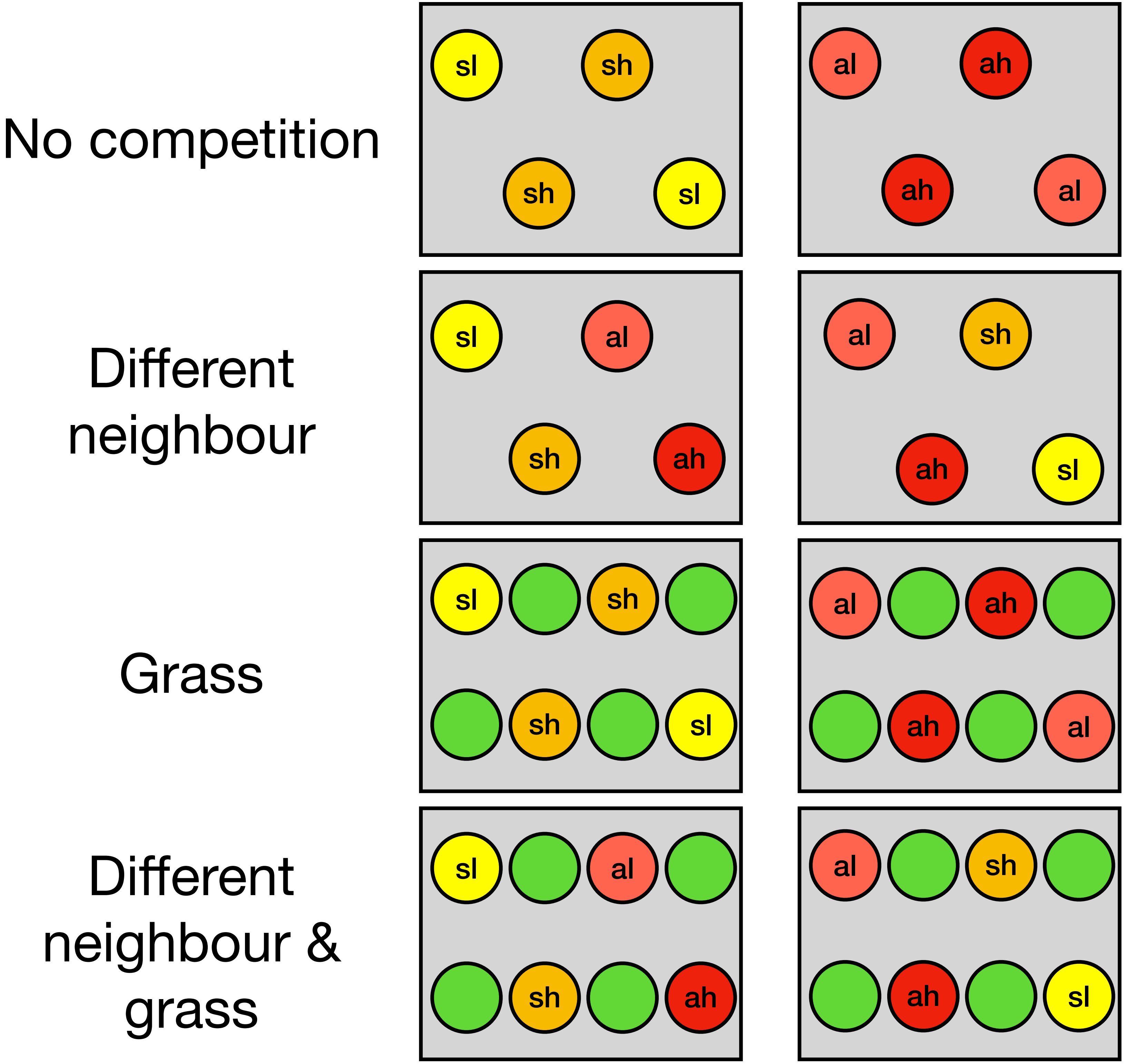


| Siblings | Mother | Father |
| --- | --- | --- |
| al & sl | MoG20-2 | low |
| ah & sh | MoG20-8 | high |
| al & sl | MoG23-8 | low |
| ah & sh | MoG23-8 | high |
| al & sl | MoK5-4 | low |

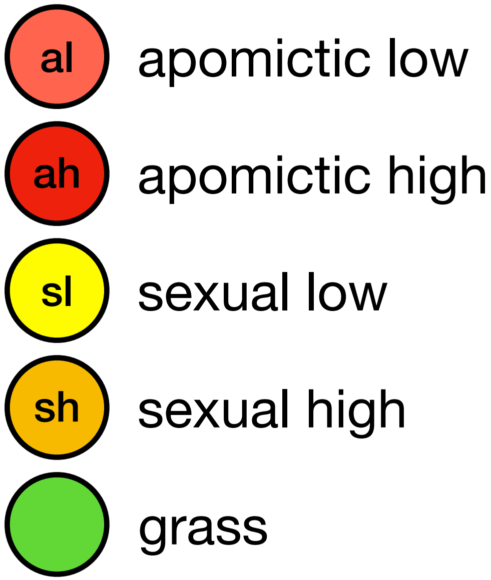
